# Signal, noise, and bias in phylogenetic inference: potential and limits to the resolution of phylogenetic trees in the phylogenomic era

**DOI:** 10.64898/2026.03.30.714540

**Authors:** Alex Dornburg, Zhuo Tony Su, Yide Jin, J. Nic Fisk, Jeffrey P. Townsend

## Abstract

Phylogenomic datasets assembled to resolve the Tree of Life now routinely span thousands of loci comprising millions of characters. Yet the persistence of incongruent topologies across such datasets reveals a fundamental truth of phylogenetics: not all data are equally informative. Here we derive analytical approaches that predict the relative impacts of phylogenetic signal, stochastic noise, and systematic bias on phylogenetic inference. We show that these three components exhibit divergent scaling properties with character sampling: signal and bias accumulate linearly, while noise accumulates nonlinearly with a concave trajectory. For some phylogenetic problems, substantial amounts of phylogenetic noise may eventually be overwhelmed by signal. For other phylogenetic problems—especially those involving deep divergences, short internodes, or constrained character-state space—the slope of signal accumulation can be so shallow that even signal from genome-scale data may never practically exceed noise. Moreover, linear accumulation of phylogenetic bias can in principle continuously overwhelm accumulation of signal at a lower slope with additional characters, regardless of dataset size. Applying our theory to empirical datasets, we show that anchored hybrid enrichment and ultraconserved element loci, like any loci, can exhibit signal that is overwhelmed by noise, and that character acquisition biases in some loci can further confound inference. Given the pervasive nature of incongruence in the phylogenomic era, our work provides a theoretical foundation for understanding the limits of inference, improving experimental design, and guiding efficient and accurate resolution of the Tree of Life.

## Introduction

Large-scale phylogenomic datasets now routinely comprise thousands of loci and millions of nucleotide characters (Singhal et al. 2021; Fedosov et al. 2024; Zuntini et al. 2024; Brownstein et al. 2025b; Stewart and Wiens 2025). Their scale has transformed phylogenetic practice by enabling the resolution of previously intractable parts of the Tree of Life (Dornburg and Near 2021; Chen et al. 2025; Torruella et al. 2025). Yet even in the phylogenomic era, major branchings of the Tree of Life remain contested, with strongly supported but conflicting topologies. Topological incongruence remains pervasive even among datasets of comparable scope, revealing a foundational problem in phylogenetics. This persistent incongruence exposes a central question in phylogenetic inference: does enough data necessarily yield a reliable tree? Indeed, we know that not all loci are equally informative (Romiguier et al. 2013; Salichos et al. 2013; Chen et al. 2015; Lewis et al. 2016; Shen et al. 2017; Martinez-Gutierrez and Aylward 2021). What remains unclear is whether some loci are deceptive. Across many empirical systems, genome-scale inferences can shift substantially depending on sampling strategy, analytical approach, or even locus selection criteria (Townsend and Lopez-Giraldez 2010; Dornburg et al. 2017a; Williams et al. 2020; Fleming et al. 2023; Steenwyk et al. 2023; Manuel et al. 2025). These persistent conflicts underscore the need to understand the conditions under which phylogenetic *signal* (differences in nucleotides or amino acids reflecting changes since common ancestry), reliably overcomes *noise* (molecular changes reflecting chance convergence or parallelism) and *bias* (molecular changes reflecting systematic convergence or parallelism).

Signal, noise, and bias are distinct components affecting the contribution of characters to phylogenetic inference, whose influence is contingent on the age of divergences under study, the temporal intervals between successive branching events, and the tempo and mode of character evolution (Ho and Jermiin 2004; Townsend et al. 2012; Su et al. 2014; Dornburg et al. 2017b; Steenwyk et al. 2023). Existing metrics for evaluating loci, such as site-rate estimates (Yang 1998; Klopfstein et al. 2017; Dornburg et al. 2019), saturation indices (Xia et al. 2003; Duchêne et al. 2022; Gosselin et al. 2022), or gene-tree congruence (Freitas et al. 2021; Singhal et al. 2021; Zhao et al. 2023), capture some important aspects of informativeness and have proven useful for identifying sources of incongruence. However, these approaches are retrospective, and do not provide a predictive theory for how signal, noise, and bias arise, accumulate, and act together within datasets. This gap in theory is particularly problematic in the phylogenomic era, where the sheer volume of data can overshadow the disproportionate contribution of a subset of loci to signal, noise, or bias, affecting overall phylogenetic inference. Without a formal theory to distinguish these components and their distinct trajectories of accumulation and interaction, phylogenomic study design and interpretation remain susceptible to unwarranted overconfidence in stochastically resolved or systematically biased inferences.

Here we build on theory from Townsend et al. (2012), Su et al. (2014), and Su and Townsend (2015) to derive a general analytical framework for prediction of the expected accumulation of signal, noise, and bias across characters and loci. We first model how signal and noise scale with character sampling, revealing their divergent accumulation dynamics and implications for inference. Explicitly accounting for rates of character change, we evaluate how lineage-specific biases in character-state usage compete with historical signal. We demonstrate that signal, noise, and bias are present and pervasive in phylogenomic datasets using (1) an avian dataset based on loci captured by anchored hybrid enrichment (Prum et al. 2015) and (2) a dataset of ultraconserved elements aimed at resolving early acanthomorph relationships (Alfaro et al. 2018). By formalizing how signal, noise, and bias scale and cumulatively interact, we expand the theoretical basis of phylogenetic experimental design providing a theory that, for the first time, anticipates the limits of accumulative character sampling on phylogenetic resolution.

### SIGNAL ACCUMULATES LINEARLY WITH CHARACTER SAMPLING

Following Townsend (2012), Su et al. (2014), Su and Townsend (Su and Townsend 2015), and Dornburg et al (2019), we consider a dataset of *n* characters and evaluate its expected utility for resolving a quartet. For each character *i* (1 ≤ *i* ≤ *n*), we allow a distinct rate of substitution on each branch of the quartet tree. Townsend et al (2012) showed that the cumulative number of characters supporting the correct subtree, whether through true synapomorphy or through homoplasy generated by parallelism or convergence, can be evaluated as

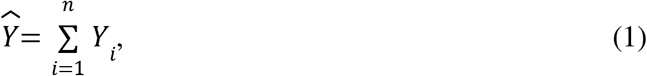

where *Y*_*i*_ = 1 if the *i*th character is in favor of the correct subtree and *Y*_*i*_ = 0 otherwise. The probability that character *i* supports the correct subtree (due to either true synapomorphy or parallelism or convergence) denoted *y*(*i*), is given by Equation 3 from Su and Townsend (2015). Because 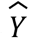 is additive across characters, its expectation increases linearly when characters are drawn from a stable distribution of evolutionary rates..

Contributions toward the correct resolution can come from true synapomorphy. However, the correct resolution can also be supported by homoplasy arising from parallelism or convergence. To measure support for correct resolution of the quartet due to actual signal only (*i.e*. “the right result for the right reason”), Townsend et al. (2012) excluded support for the correct quartet subtree due to homoplasy (*i.e*. “the right result for the wrong reason”). The contribution of each character *i* can be reduced by the probability that each character represents homoplasy (Appendix), leaving an expected number of characters that exhibit true synapomorphy of 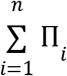. The amount of signal remains a simple sum across characters in a given alignment, thereby accumulating linearly with increased character sampling (**Fig. 1A**).

**Figure 1.**
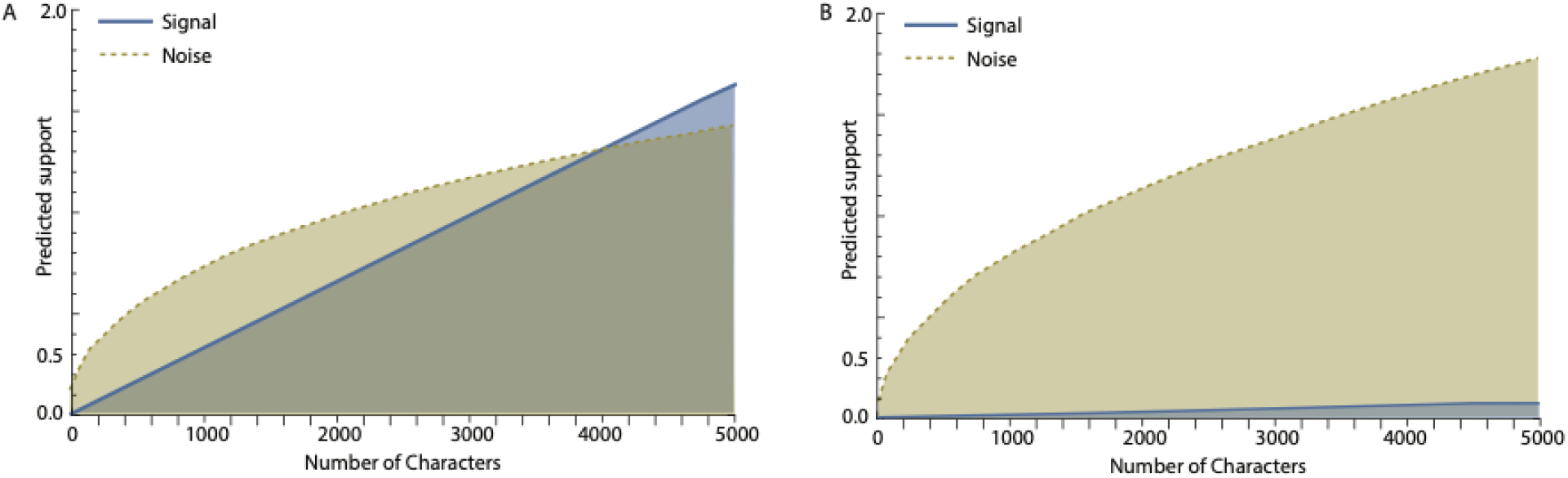
Expected levels of signal and noise on a quartet tree with subtending branches of lengths of length *T* = 1 as a function of evenly sampling of up to 5000 nucleotide sequence characters, of which 1/33 evolve at a near-optimal rate (0.25), 1/33 evolve at four times the optimal rate (1), and 31/33 evolve at a very slow rate (1 × 10^−5^), and for which base frequencies are all equal to 0.25. (**A**) Signal and noise expected when *t*_0_ = 0.1. (**B**) Signal and noise expected when the internode is 20-fold shorter (*t*_0_ = 0.005) than that used for panel A.

### NOISE ACCUMULATES NONLINEARLY WITH CONCAVE SLOPE

A quartet tree has three possible tip-labeled subtrees. Only one of the subtrees, denoted as τ_3_, matches the actual quartet topology, while the other two subtrees, denoted as τ_1_ and τ_2_, are incorrect (*c.f*. Figure 1 in Townsend et al. 2012). For each of the two incorrect quartet subtrees τ_1_ and τ_2_, we respectively define 2*n* indicator random variables *X*_1,*i*_ and *X*_2,*i*_ over the domain 1 ≤ *i* ≤ *n. X*_1,*i*_ = 1 if the *i-*th character is in favor of the incorrect subtree τ_1_, otherwise *X*_1,*i*_ = 0. The probability of the *i-*th character character supporting the incorrect subtree τ_1_ can be represented by *x*_1_ (*i*) (Equation 4 in Su and Townsend (2015)). Assuming no bias in character states and independence of *X*_1,*i*_ across all the *n* characters, the cumulative number of homoplasious characters in the data set that are in favor of the incorrect topology τ_1_ can be evaluated as 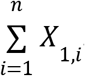. The *n* variables *X*_2,*i*_ are defined similarly so as to indicate the number favoring the incorrect subtree τ_2_, 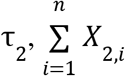. The probability of the *i-*th character supporting τ_2_ is *x*_2_ (*i*), and is given by Equation 5 in Su and Townsend (2015).

Noise can also yield support for the correct topological resolution as a result of parallelism or convergence. The *n* variables *X*_3,*i*_ can be defined analogously to *X*_1,*i*_ and *X*_2,*i*_, so that the number of characters favoring the correct subtree τ_3_ due to noise is 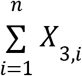. The corresponding probability that the *i*th character supports the correct subtree due to parallelism or convergence but not due to true synapomorphy can be quantified by evaluating the probability that the *i*th character supports the correct subtree (due to either true synapomorphy or parallelism or convergence) and subtracting the probability that the *i*th character supports the correct subtree due to true synapomorphy, so that *x*_3_ (*i*) = *y*(*i*) − Π(*i*).

To measure support for incorrect resolution of the quartet, we can contrast the number of homoplasious characters in favor of the correct subtree with the number of homoplasious characters in favor of the more supported of the two incorrect subtrees. This discounted contingent net support for incorrect quartet resolution without bias

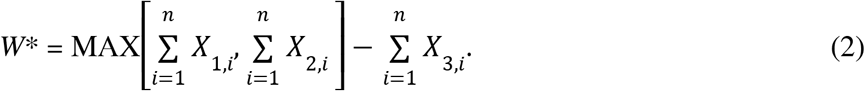

Intuitively, the maximum of two sums of identically distributed random variables subtracting a single identically distributed random variable is unlikely to be linear. Indeed, because of a large-magnitude square-root term arising from the “random walk” with increased i.i.d. character sampling of the competing incorrect trees and correct tree (**Appendix 1**), the expected noise is nonlinear:

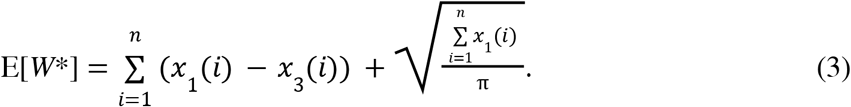

To better understand this function, let’s consider some extremes. If there is no true signal of the phylogenetic relationship at all (e.g. if the internode of interest is a hard polytomy), then Π(*i*) = 0 and *x*_3_ (*i*) = *x*_1_ (*i*), such that

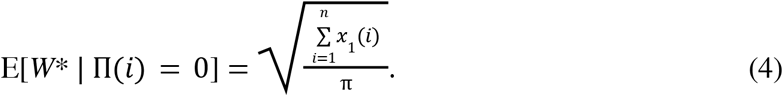

This square-root function is an intuitive result of the competition of three i.i.d. unbiased random walks, each one conferring support to one of the three potential resolutions of a phylogenetic quartet. The nonlinearity of E[*W\**] provides a concave slope with respect to data set size (**Fig. 1A**). Relatively speaking, phylogenetic noise is always more problematic to phylogenetic inference when the size of a data set is small; signal tends to overwhelm noise as the size of the dataset increases. This result is a mathematical basis for the conventional wisdom that signal will eventually overcome noise, and that with enough sequence data one can always “sample one’s way out of …”.

Unfortunately, this conventional wisdom is not always correct. Depending in particular on the shortness of the internodes, their depth in the phylogeny, and their rate of evolution, the slope of signal can be arbitrarily low (**Fig. 1B**). Furthermore, at very high levels of sampling, the square-root term in Eq. 4 eventually diminishes in impact compared to the linear term, so that noise itself becomes increasingly linear. Even if the linear slope of signal is greater than the asymptotically linear slope of noise, the finite number of characters to sample can fail to reach the “inevitable” crossing-point, and render a phylogenetic problem insoluble.

### BIAS ADDS LINEARLY TO NOISE WITH CHARACTER SAMPLING

The cumulative number of characters in support of the incorrect subtrees (*W*) arises from homoplasy stemming from parallelism or convergence. When parallelism or convergence manifest as heterogeneous base composition among species driven by lineage-specific processes such as biased gene conversion (Lartillot 2013; Bolívar et al. 2019; Rousselle et al. 2019), one component of the noise can be differentiated and termed phylogenetic bias. Phylogenetic bias misleads phylogenetic inference as a consequence of distantly related lineages systematically converging in character states because of shared biases in character state frequencies. Recent research has demonstrated that such convergence presents a particularly notable challenge within large phylogenomic-scale datasets (Dornburg et al. 2017a; de Moya et al. 2021; Höhna et al. 2025). Quantifying the potential impact of phylogenetic bias therefore represents a fundamental challenge that must be addressed to guide data collection and conduct robust phylogenetic inference.

To evaluate the relative contribution of phylogenetic bias, noise can be evaluated for any evolutionary model and corresponding character rate vector, with and without lineage-specific differences in state frequencies across characters, such as nucleotide or amino-acid base composition. Noise that can be attributed to heterogeneity of character state frequencies across lineages can be isolated by taking the difference between the total noise *W* (with composition bias) and the total noise without composition bias *W*\*. Thus, phylogenetic bias

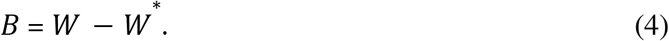

When sampling characters randomly from any fixed dataset, bias accrues linearly (**Fig. 2**). The linear slope of the phylogenetic bias as characters are sampled can be lower than the linear slope of signal (**Fig. 2A**)—or higher (**Fig. 2B**). High slopes of phylogenetic bias are expected to manifest on an evolutionary timescale: there are numerous cases of GC bias in specific lineages (Romiguier et al. 2016; Bossert et al. 2017; Dornburg et al. 2017a; Rousselle et al. 2019; White and Braun 2019), and essentially limitless potential for arbitrarily short internodes that depauperate the slope of signal (Dornburg et al. 2019). This potentially higher slope of phylogenetic bias with character sampling is why the conventional wisdom that signal will eventually overcome other topological estimation errors—and the idea that with enough sequence data one can always “sample one’s way out of …”—is not only problematic in the ubiquitous circumstance of finite numbers of characters, but false even if limitless characters were available.

**Figure 2.**
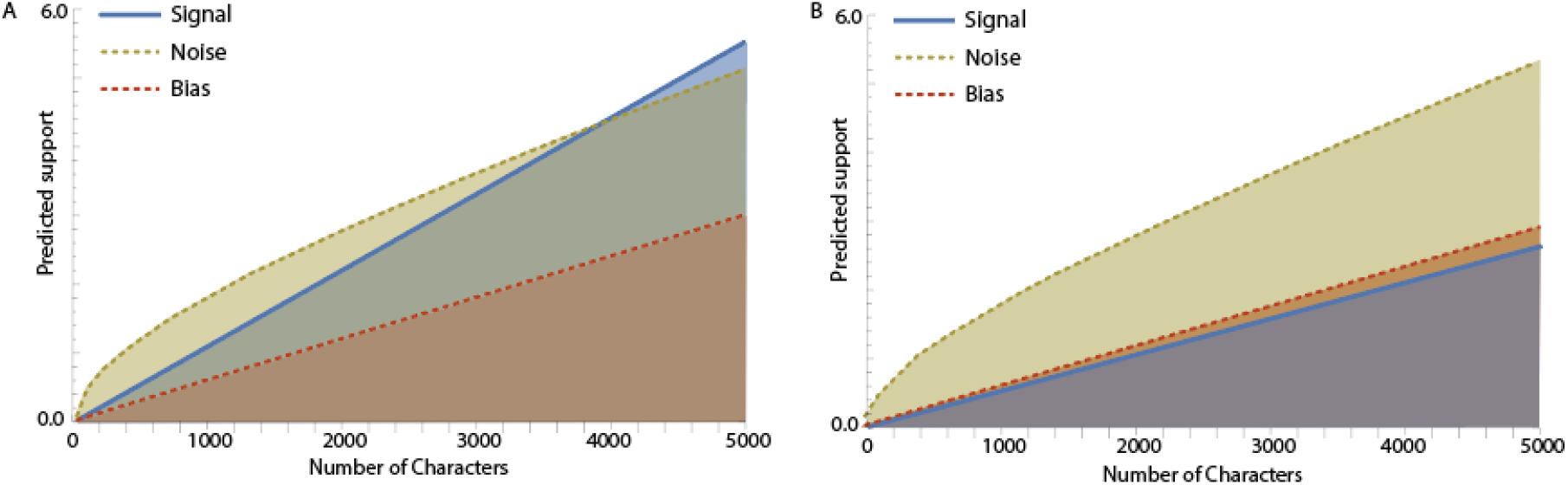
Expected signal, noise, and phylogenetic bias on a quartet tree with subtending branches of length *T* = 1 as a function of even sampling of up to 5000 nucleotide sequence characters, (**A**) when 6/70 of sites evolve at a near-optimal rate (0.25), 8/70 evolve at four times that near-optimal rate (1.0), and 55/70 evolve at an extremely slow rate (1 × 10^−5^), and (**B**) when all sites evolve at twice the rates as in panel A.

### IT GETS EVEN WORSE…

*When combined with lineage-specific biases toward a subset of character states, fast rates of sequence evolution, longer divergence times, and lineage-specific rate heterogeneity can amplify the impact of phylogenetic bias*.

Not only can high rates of character change throughout a phylogeny increase the slope of phylogenetic bias with character sampling, rates of character change can vary heterogeneously among lineages (Field et al.; Soltis et al. 2002; Dornburg et al. 2012; Beaulieu et al. 2015; Berv and Field 2018; Vankan et al. 2022; Berv et al. 2024). A worst-case scenario of lineage-specific rate heterogeneity for phylogenetic inference is that of long-branch attraction, in which non-sister lineages share an elevated rate of evolution (Hendy and Penny 1989; Huelsenbeck 1997; Philippe et al. 2005; Su and Townsend 2015), a scenario that markedly increases phylogenetic noise (Fig. 3A–B; Su and Townsend 2015). The presence of lineage-specific rates can also increase the slope of phylogenetic bias with character sampling. Even as a consequence of long-branch attraction, however, persistent noise may eventually be overcome by linear accumulation of signal (e.g. **Fig. 3A**). With phylogenetic bias, inference of the true branching structure of a quartet in which two non-sister taxa branches have longer branch lengths can be subject not only to this well-known long-branch attraction effect due to noise and saturation leading to spurious, stochastic parallelism and convergence (Su and Townsend 2015), but also to a nefarious, higher linear slope of phylogenetic bias with character sampling (**Fig. 3C–D**). In short, the linear phylogenetic bias that cannot be overcome by additional sampling becomes amplified.

**Figure 3.**
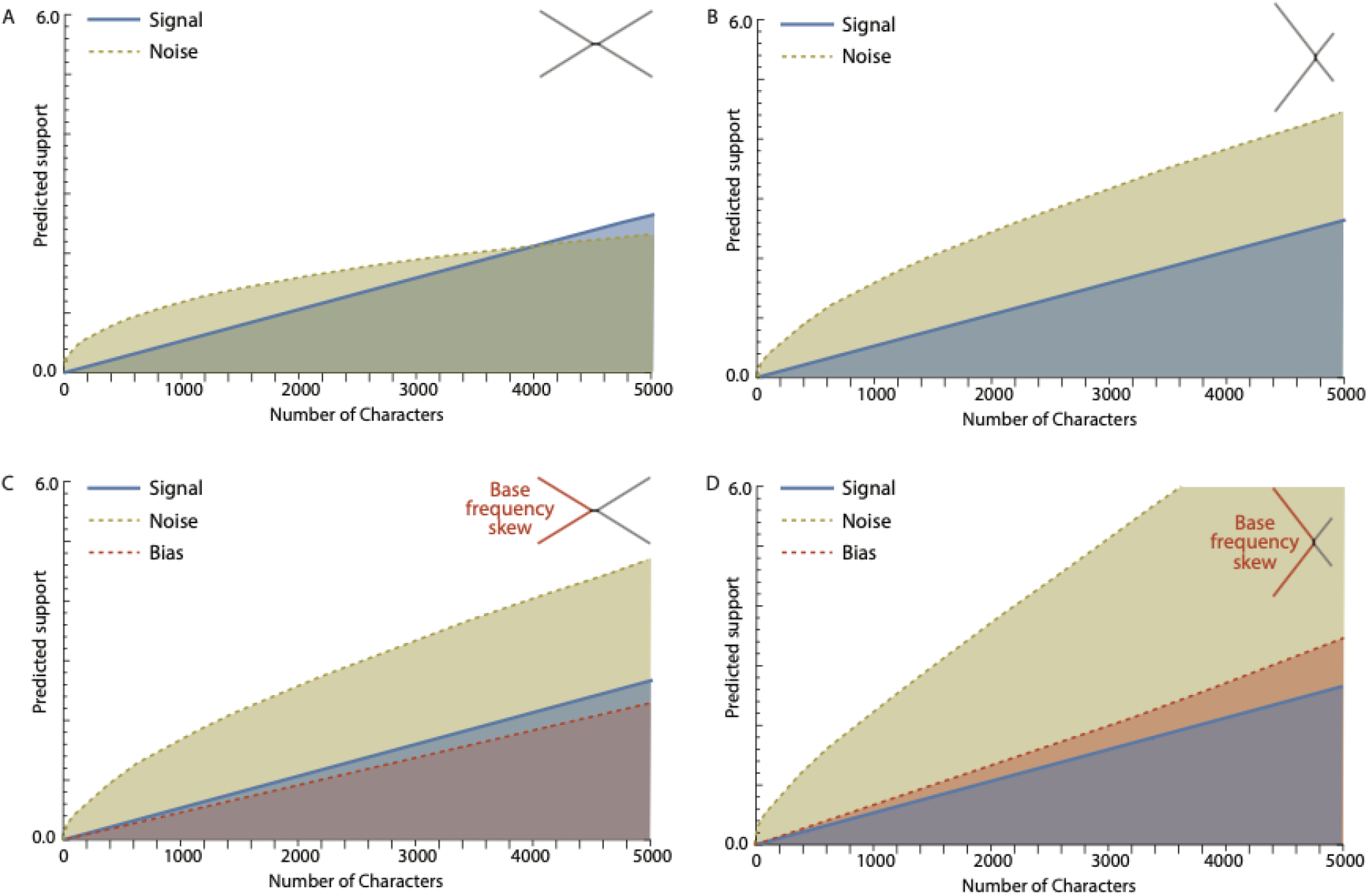
Expected value of signal (solid line), noise (dashed line), and bias (dotted line) of a hypothetical data set for resolving a quartet phylogeny, versus the number of nucleotide characters in the data set, (**A**) when 6/70 of sites evolve at a near-optimal rate (0.25), 8/70 evolve at four times that near-optimal rate (1.0), and 55/70 evolve at an extremely slow rate (1 × 10^−5^), according to the Jukes and Cantor model. The branch lengths of the quartet tree in terms of evolutionary time are 1 unit time for the four terminal branches and 0.1 unit time for the internode; (**B**) when rates are the same as in panel A but the branch lengths of the quartet tree in evolutionary time are 1.3 time units for the two non-sister quartet branches, 0.7 for the sister quartet branches, and 0.1 for the deep internode; (**C**) when all parameters are the same as in panel A but the nucleotide base frequencies for two non-sister quartet branches are *π*_*T*_ = 0.34, *π*_*C*_ = 0.16, *π*_*A*_ = 0.34, and *π*_*G*_ = 0.16, and nucleotide base frequencies for the short deep internode are *π*_*0*,*T*_ = 0.30, *π*_*0*,*C*_ = 0.20, *π*_*0*,*A*_ = 0.30, and *π*_*0*,*G*_ = 0.20; and (**D**) when nucleotide base composition frequencies are skewed as in panel C, and quartet branch lengths are unequal—as in panel B, with the two longer quartet branches being the two non-sister branches that are equivalently frequency-skewed.

### CHARACTER-ACQUISITION BIAS IS NOT PHYLOGENETIC BIAS

Deviations from base compositional stationarity are a common source of model violation in phylogenetics and can substantially influence tree inference (Romiguier and Roux 2017; Jermiin et al. 2020; Kapli et al. 2020; Rota et al. 2022; Naser-Khdour et al. 2026). Deviations are often treated as a singular issue of compositional heterogeneity, but in practice, they encompass at least two distinct and underappreciated analytic challenges. First, character-acquisition biases, such as skewed nucleotide or codon usage, can reduce the effective dimensionality of the accessible state space, thereby compressing phylogenetic signal and amplifying the impact of stochastic noise (Townsend et al. 2012; Dornburg et al. 2019). Second, heterogeneity in character biases among lineages can introduce systematic phylogenetic bias, favoring incorrect topologies through convergent patterns of character states (Bossert et al. 2017; Dornburg et al. 2017a; de Moya et al. 2021). Non-stationarity is frequently flagged as a confounding factor. However, the entanglement of these two sources of error, noise inflation and directional bias, is rarely disentangled in empirical studies.

To illustrate these issues, we evaluated the component contributions of signal, noise, and phylogenetic bias in a widely used dataset of anchored hybrid enrichment loci (Prum et al. 2015), generated to resolve deep relationships in the avian tree of life. We focused on the placement of the Hoatzin (*Opisthocomus hoazin*), a lineage whose phylogenetic position has remained notably unstable across recent analyses (Jarvis et al. 2014; Prum et al. 2015; Stiller et al. 2024). Using the divergence times, site-specific rates, and sequence alignments reported by Prum et al. (2015), we estimated the relative expected contributions of phylogenetic signal, stochastic noise, and potential bias for the internode relevant to the placement of the Hoatzin. Our results reveal a striking pattern: for this internode, every locus is expected to contribute more noise than signal (**Fig. 4A–B**), whereas fewer than 5% exhibit bias exceeding signal (**Fig. 4C**). This near absence of phylogenetic bias is in accord with the relatively low-variance distribution of elevated AT content across the lineages surrounding the focal internode (**Fig. 4D**). This low variance in AT content among lineages limits any directional tendency toward an incorrect resolution. In this case, incongruence is driven far more by noise than by bias. This example illustrates an important facet of phylogenetic study design: a uniform bias in the evolution of characters does not necessarily result in phylogenetic bias.

**Figure 4.**
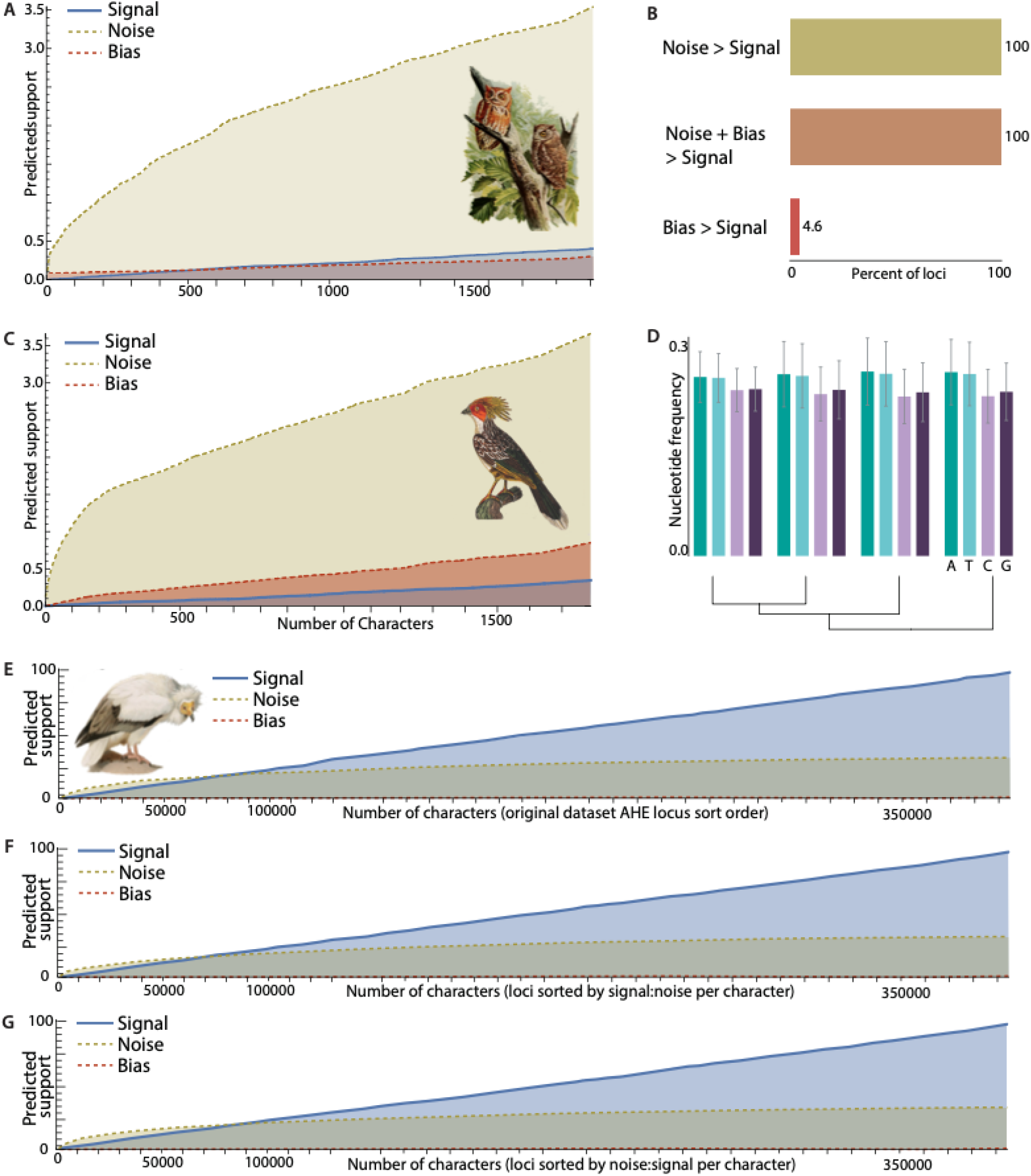
Analysis of predicted phylogenetic signal, stochastic noise, and phylogenetic bias regarding placement of the Hoatzin (*Opisthocomus hoazin*) in *n* = 259 anchored hybrid enrichment loci. (**A**) An anchored hybrid enrichment locus (#229) that provides nearly equivalent amounts of signal and bias, and considerably greater noise. (**B**) Percent of loci that as individual loci provide greater noise than signal, greater noise plus bias than signal, and greater bias than signal for the internode that the Hoatzin subtends. (**C**) An anchored hybrid enrichment locus (#24) that provides greater bias than signal. (**D**) uniformity of elevated AT content among lineages surrounding the internode for the four taxa subtending the ancestor of the hoatzin. (**E**) Predicted signal, noise, and bias for the concatenation of all anchored hybrid enrichment loci. (**F**) Predicted signal, noise, and bias for the concatenation of all anchored hybrid enrichment loci when sorted by their ratios of signal:noise per character. (**G**) Predicted signal, noise, and bias for the concatenation of all anchored hybrid enrichment loci when sorted by the their ratios of noise:signal per character.

When base-frequency proportions are elevated among all lineages, character-acquisition bias has an effect similar to reducing the dimensionality of the character-state space. In such a case, stochastic noise is amplified (Townsend et al. 2012; Su et al. 2014; Dornburg et al. 2019). This interaction between rate of evolution and character state space can be exacerbated by the combination of deep time scales and short internodes (Townsend 2007; Townsend et al. 2012). In the case of the unstable placement of the Hoatzin (Jarvis et al. 2014; Prum et al. 2015; Cracraft 2022; Stiller et al. 2024), for example, signal is expected to exceed noise only after tens of thousands of characters have been sampled (**Fig. 4E**), regardless of the order in which loci are added (**Fig. 4E–G)**.

These considerations on signal accumulation extend beyond the placement of the hoatzin and reflect broader biologically driven biases in character acquisition. A well-established example is codon usage bias, in which synonymous codons are employed non-randomly because of a combination of mutational pressures, translational selection shaped by tRNA abundance, and selection on mRNA stability and structure (Quax et al. 2015; Zhou et al. 2016; Frumkin et al. 2018; Hanson and Coller 2018; Wu et al. 2024). Such biases restrict the range of accessible substitutions in protein-coding regions, effectively reducing the dimensionality of the character-state space and thereby increasing the relative accumulation of noise and bias while limiting phylogenetically informative variation. These molecular constraints can reinforce non-stationary patterns of evolution, particularly at third codon positions, impeding the recovery of accurate topologies from individual loci and even from genome-scale datasets. Accordingly, more than a decade of work has emphasized the importance of detecting and modeling deviations from compositional stationarity in phylogenetics (Betancur-R et al. 2013; Lartillot 2013; Dornburg et al. 2017a; Parker et al. 2019; White and Braun 2019). Our results provide a complementary perspective: by predicting how signal, noise, and bias accumulate under such constraints, it enables more efficient, targeted data collection as well as facilitating a rigorous basis for understanding why particular loci, or datasets enriched with similar loci, may contribute disproportionately to phylogenetic conflict or congruence.

### NOISE AND BIAS ARE PERVASIVE IN EMPIRICAL DATASETS

Most phylogeneticists working with empirical datasets encounter inconsistency, low support, or conflicting topologies, even when utilizing high-volume sequencing techniques and sophisticated models of molecular evolution (Philippe et al. 2011; Reddy et al. 2017; Young and Gillung 2020; Foley et al. 2023). It is true that technological advances and model refinement have certainly improved phylogenetic resolution (Nagy and Szöllősi 2017; Wang et al. 2025). However, they alone cannot circumvent a core problem in phylogenetics. Not all data are equally informative, even when collected and analyzed with modern methods. Phylogenomic datasets differ in rate, composition, and evolutionary histories, and these factors shape the relative contributions of signal, noise, and bias in ways that are difficult to anticipate without formal evaluation.

To demonstrate this issue, we analyzed a published ultraconserved element (UCE) dataset of >1001 loci used to study the evolutionary history of Acanthomorpha (Alfaro et al. 2018), focusing on the phylogenetic placement of sleepers (Kurtidae), a lineage whose position has been incongruent between studies (Alfaro et al. 2018; Kuang et al. 2018; Ghezelayagh et al. 2022), including attempts to resolve the early evolutionary history of Gobiiformes (Dornburg and Near 2021; Near and Thacker 2024). UCEs are commonly used in phylogenomics (Chakrabarty et al. 2017; Brownstein et al. 2024, 2025a) and often presumed to be robust phylogenetic markers because of their conserved central regions flanked by more rapidly evolving sites that can be informative at different time depths (Faircloth et al. 2012; Zhang et al. 2019). Consistent with this expectation, most loci depict an initial accumulation of signal across the first few hundred characters, followed by a plateau within the ultraconserved core, then a subsequent increase in the adjacent flanking element (**Fig. 5A**). However, across all 1001 loci, levels of noise exceeded signal (**Fig. 5B**). In a small number of cases, signal was overwhelmed by the combination of noise and bias (**Fig. 5C**). As in the case of the AHE dataset, phylogenetic bias was low as all four branches of the quartet showed parallel enrichments of Adenine and Thymine (**Fig. 5D**). These results highlight that widely used genomic markers like UCEs are susceptible to the same forces that complicate single-gene phylogenetics, especially when evolutionary history is complex or internodes are short relative to subtending branches. In the case of these data, high noise within the individual loci can explain the empirical discordance observed between gene-tree species tree analyses and concatenation (McCraney et al. 2025).

**Figure 5.**
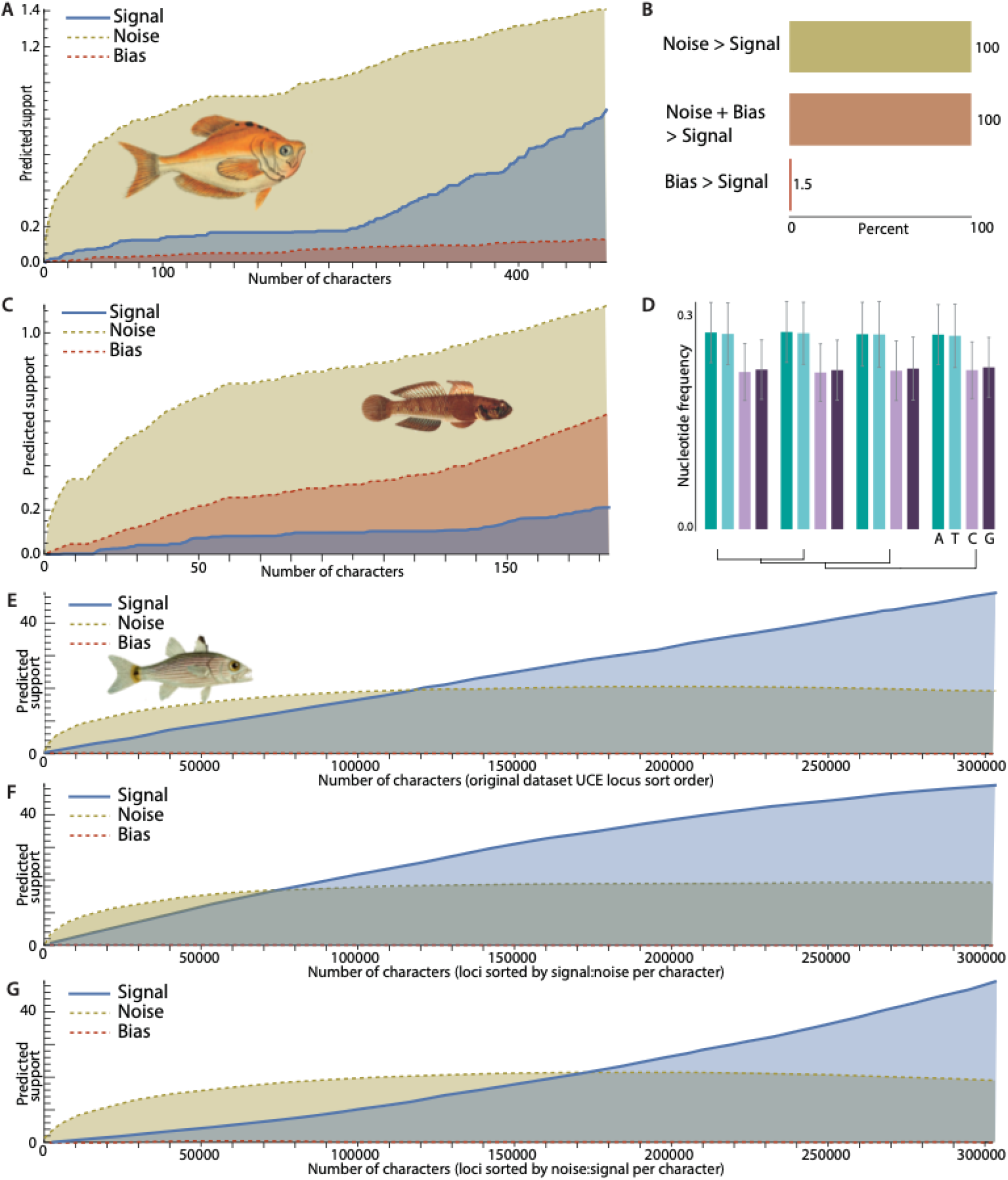
Analysis of predicted phylogenetic signal, stochastic noise, and bias regarding the evolutionary history of sleepers (Kurtidae) for *n* = 1001 ultraconserved element loci. (**A**) An ultraconserved element locus (121) that provides greater predicted noise than signal, and minimal predicted bias. (**B**) Percent of loci that as individual loci provide greater noise than signal, greater noise plus bias than signal, and greater bias than signal for the internode associated with sleepers. (**C**) An ultraconserved element locus (511) that provides greater predicted noise than signal, and substantial predicted bias. (**D**) Uniformity of elevated AT content among lineages surrounding the internode for the four taxa subtending the ancestor of the sleepers. (**E**) Predicted signal, noise, and bias for the concatenation of all 1001 ultraconserved element loci in the original dataset order. (**F**) Predicted signal, noise, and bias for the concatenation of all anchored hybrid enrichment loci when sorted by their ratios of signal:noise per character. (**G**) Predicted signal, noise, and bias for the concatenation of all ultraconserved element loci when sorted by the their ratios of noise:signal per character.

Our results provide additional insight into the observation that even concatenated UCE datasets can underperform at resolving certain nodes (Borowiec et al. 2025). Specifically, signal may only surpass noise after tens of thousands of characters have been sampled (**Fig. 5E**)—and that the order in which loci are added can critically shape this trajectory (**Fig. 5E–G)**. These findings further underscore a critical—yet often overlooked—aspect of phylogenomic study design. A nontrivial fraction of loci may impede inference more than they contribute to it, and would be better off left unsampled.

This point is readily demonstrated by altering the specific sequence in which loci are added in an cumulative signal, noise, and bias plot. Our UCE-based analysis shows that the order of sampling can dramatically alter the expected amount of data required for signal to surpass noise (**Fig. 5E–G**). For example, the UCE dataset as assembled in the originally reported locus order (**Fig 5E**) requires approximately 110,000 characters before signal begins to outpace noise. This cost could have been avoided with more strategic data selection (**Fig. 5F**) or made worse through inadvertent selection of the loci with the highest noise:signal values (**Fig. 5G**). Moreover, the trajectory of signal of the sorted datasets demonstrates a marked convexity (**Fig. 5F**) or concavity (**Fig. 5G**) depending on locus ordering. The concavity reflects the early accumulation of relatively informative loci followed by a subsequent addition of relatively noise-ridden loci. The inverse is true when prioritizing loci with high noise:signal per character. This worst-case scenario demonstrates how early inclusion of noisy loci can dramatically inflate the data requirements for accurate inference. Collectively, our results reinforce that data addition is not equivalent across loci, manifesting the central role of experimental design in achieving efficient and reliable phylogenetic resolution. Since our framework calculates expected noise continuously across characters, it captures how early inclusion of noisy loci can delay the accumulation of higher signal to noise levels per character. This approach not only clarifies why some phylogenetic questions remain stubbornly unresolved despite large datasets, but also offers a path forward: one where signal, noise, and bias can be explicitly modeled to guide informed, efficient sampling strategies.

## Discussion

Here we have derived an analytical framework that quantifies how phylogenetic signal, noise, and bias accumulate as character sampling increases. We demonstrate that signal and bias accumulate linearly, whereas noise accumulates nonlinearly with a concave trajectory. This simple yet powerful distinction provides fundamental insight into phylogenetic experimental design: Noise can easily initially exceed signal. However, its growth rate slows, in principle allowing signal to eventually dominate. To the misfortune of inference, that “inevitable” domination of signal over noise may never come about. It is always possible for an internode of currently unknown branch length to be in actuality sufficiently short such that even genome-scale sampling would never provide sufficient information to overcome noise.

Bias presents an even more diabolical hazard. Because it contributes linearly to noise in supporting incorrect resolution, bias can in some cases drive resolution towards an incorrect topology faster than signal can accumulate. These theoretical expectations were borne out in two empirical datasets, where even data types commonly viewed as reliable—anchored hybrid enrichment loci and ultraconserved elements—contain substantial proportions of loci with noise exceeding signal, or with non-negligible bias.

This work provides a quantitative resolution to a foundational debate in phylogenetics: under what conditions does increasing character sampling overcome incongruence? One view has long held that with enough data, any phylogenetic problem can be resolved (Wortley et al. 2005; Spinks et al. 2009). Others have emphasized that certain questions remain highly challenging, if not intractable, due to noise or bias (Philippe et al. 2011; Haber 2025). Our results reconcile these views showing that these positions are not contradictory, but conditional. Whether signal eventually dominates depends not just on the quantity of data, but on the slope of signal accumulation relative to the nonlinear rise of noise and the linear compounding of bias. This formalization offers new insight into when adding data is expected to improve inference and when it will fail, and provides a predictive foundation for phylogenetic study design. Decisions about the sampling of loci, taxa, and characters can be evaluated in light of expected information trajectories rather than by raw intuition.

One important caveat of this framework is that it assumes approximate uniformity of branch lengths and therefore does not directly account for long branch attraction (LBA). LBA occurs when substantial branch length or rate asymmetries increase the probability that causes distantly related taxa to cluster incorrectly due to shared noise (McTavish et al. 2015; convergence or parallelism; Susko 2015; Ontano et al. 2021; Szánthó et al. 2023). Within our framework, these effects are most naturally interpreted as an elevation of noise, consistent with previous theory showing that long, uneven branches increase the probability of stochastic convergence (Su et al. 2014). Under conditions of extreme branch-length heterogeneity, noise is expected to accumulate more steeply, shifting the threshold at which signal can overwhelm noise. The current formulation does not explicitly model such asymmetry. However, it could be extended using established theoretical results on branch-length heterogeneity. Future applications of this framework to empirical datasets with pronounced branch-length disparities could apply this theory to derive a minimal expectation of noise.

It is important to emphasize that our theoretical framework predicts *expected* information content based on rates of character change rather than realized informativeness in any particular dataset. Once datasets have been assembled, these study design approaches should be complemented by approaches that assess empirical informativeness. Metrics such as gene/site concordance factors, for example, provide critical validation of these predictions in practice (Minh et al. 2020; Mo et al. 2023; Lanfear and Hahn 2024). Importantly, model choice and partition selection during inference can have a profound impact on estimated topologies (Susko and Roger 2021; Al Jewari and Baldauf 2023; McCarthy et al. 2023; Cai 2024) the factors that play into these empirical assessments have limited influence on the rate-derived signal, noise, and bias trajectories we calculate here (Su et al. 2014). Selection of data that maximizes power and minimizes error is foundational to obtaining robust parameter estimates in any field, including phylogenetics (Dornburg et al. 2017b; Fleming et al. 2023; Grigoriadis et al. 2024). By identifying sets of loci or genomic partitions whose collection will maximize signal while minimizing bias and noise, this theory provides a means for justification of proposed data collection and can guide an expectation of success before data are collected and when funding decisions are made for phylogenetic studies. Operating upstream of novel research study data collection, model selection and inference, this conception of the interplay of phylogenetic signal, noise, and bias provides a critical lens through which the design and analysis of phylogenetic studies can be optimized in the genomic era.

## Acknowledgements

We thank Steven Wang for executing preliminary analyses on early datasets.

## Data Availability Statement

Mathematica notebooks and input files used in these analyses are available on Zenodo (DOI: 10.5281/zenodo.19239631; https://zenodo.org/records/19239632). R code to replicate these analyses is available from github: https://github.com/yide0202/SignalNoiseBias_R.

## References

Alfaro M.E., Faircloth B.C., Harrington R.C., Sorenson L., Friedman M., Thacker C.E., Oliveros C.H., Černý D., Near T.J. 2018. Explosive diversification of marine fishes at the Cretaceous-Palaeogene boundary. Nat Ecol Evol. 2:688–696.

Al Jewari C., Baldauf S.L. 2023. Conflict over the Eukaryote Root Resides in Strong Outliers, Mosaics and Missing Data Sensitivity of Site-Specific (CAT) Mixture Models. Syst Biol. 72:1–16.

Beaulieu J.M., O’Meara B.C., Crane P., Donoghue M.J. 2015. Heterogeneous Rates of Molecular Evolution and Diversification Could Explain the Triassic Age Estimate for Angiosperms. Syst. Biol. 64:869–878.

Berv J.S., Field D.J. 2018. Genomic Signature of an Avian Lilliput Effect across the K-Pg Extinction. Syst. Biol. 67:1–13.

Berv J.S., Singhal S., Field D.J., Walker-Hale N., McHugh S.W., Shipley J.R., Miller E.T., Kimball R.T., Braun E.L., Dornburg A., Parins-Fukuchi C.T., Prum R.O., Winger B.M., Friedman M., Smith S.A. 2024. Genome and life-history evolution link bird diversification to the end-Cretaceous mass extinction. Sci Adv. 10:eadp0114.

Betancur-R R., Li C., Munroe T.A., Ballesteros J.A., Ortí G. 2013. Addressing gene tree discordance and non-stationarity to resolve a multi-locus phylogeny of the flatfishes (Teleostei: Pleuronectiformes). Syst. Biol. 62:763–785.

Bolívar P., Guéguen L., Duret L., Ellegren H., Mugal C.F. 2019. GC-biased gene conversion conceals the prediction of the nearly neutral theory in avian genomes. Genome Biol. 20:5.

Borowiec M.L., Zhang Y.M., Neves K., Ramalho M.O., Fisher B.L., Lucky A., Moreau C.S. 2025. Evaluating UCE Data Adequacy and Integrating Uncertainty in a Comprehensive Phylogeny of Ants. Syst Biol.

Bossert S., Murray E.A., Blaimer B.B., Danforth B.N. 2017. The impact of GC bias on phylogenetic accuracy using targeted enrichment phylogenomic data. Mol. Phylogenet. Evol. 111:149–157.

Brownstein C.D., Dornburg A., Near T.J. 2025a. Cenozoic evolutionary history obscures the Mesozoic origins of acanthopterygian fishes. Evolution. 79:922–934.

Brownstein C.D., Harrington R.C., Alencar L.R.V., Bellwood D.R., Choat J.H., Rocha L.A., Wainwright P.C., Tavera J., Burress E.D., Muñoz M.M., Cowman P.F., Near T.J. 2025b. Phylogenomics establishes an Early Miocene reconstruction of reef vertebrate diversity. Sci Adv. 11:eadu6149.

Brownstein C.D., Zapfe K.L., Lott S., Harrington R., Ghezelayagh A., Dornburg A., Near T.J. 2024. Synergistic innovations enabled the radiation of anglerfishes in the deep open ocean. Curr Biol. 34:2541–2550.e4.

Cai C. 2024. Ant backbone phylogeny resolved by modelling compositional heterogeneity among sites in genomic data. Commun Biol. 7:106.

Chakrabarty P., Faircloth B.C., Alda F., Ludt W.B., Mcmahan C.D., Near T.J., Dornburg A., Albert J.S., Arroyave J., Stiassny M.L.J., Sorenson L., Alfaro M.E. 2017. Phylogenomic Systematics of Ostariophysan Fishes: Ultraconserved Elements Support the Surprising Non-Monophyly of Characiformes. Syst Biol. 66:881–895.

Chen M.-Y., Liang D., Zhang P. 2015. Selecting Question-Specific Genes to Reduce Incongruence in Phylogenomics: A Case Study of Jawed Vertebrate Backbone Phylogeny. Syst. Biol. 64:1104–1120.

Chen Z., Baeza J.A., Chen C., Gonzalez M.T., González V.L., Greve C., Kocot K.M., Arbizu P.M., Moles J., Schell T., Schwabe E., Sun J., Wong N.L.W.S., Yap-Chiongco M., Sigwart J.D. 2025. A genome-based phylogeny for Mollusca is concordant with fossils and morphology. Science. 387:1001–1007.

Cracraft J. 2022. The Hoatzin. Curr Biol. 32:R1068–R1069.

Dornburg A., Brandley M.C., McGowen M.R., Near T.J. 2012. Relaxed clocks and inferences of heterogeneous patterns of nucleotide substitution and divergence time estimates across whales and dolphins (Mammalia: Cetacea). Mol. Biol. Evol. 29:721–736.

Dornburg A., Near T.J. 2021. The emerging phylogenetic perspective on the evolution of actinopterygian fishes. Annu. Rev. Ecol. Evol. Syst. 52:427–452.

Dornburg A., Su Z., Townsend J.P. 2019. Optimal Rates for Phylogenetic Inference and Experimental Design in the Era of Genome-Scale Data Sets. Syst. Biol. 68:145–156.

Dornburg A., Townsend J.P., Brooks W., Spriggs E., Eytan R.I., Moore J.A., Wainwright P.C., Lemmon A., Lemmon E.M., Near T.J. 2017a. New insights on the sister lineage of percomorph fishes with an anchored hybrid enrichment dataset. Mol. Phylogenet. Evol. 110:27–38.

Dornburg A., Townsend J.P., Wang Z. 2017b. Maximizing Power in Phylogenetics and Phylogenomics: A Perspective Illuminated by Fungal Big Data. Adv. Genet. 100:1–47.

Duchêne D.A., Mather N., Van Der Wal C., Ho S.Y.W. 2022. Excluding Loci With Substitution Saturation Improves Inferences From Phylogenomic Data. Syst Biol. 71:676–689.

Faircloth B.C., McCormack J.E., Crawford N.G., Harvey M.G., Brumfield R.T., Glenn T.C. 2012. Ultraconserved elements anchor thousands of genetic markers spanning multiple evolutionary timescales. Syst. Biol. 61:717–726.

Fedosov A.E., Zaharias P., Lemarcis T., Modica M.V., Holford M., Oliverio M., Kantor Y.I., Puillandre N. 2024. Phylogenomics of Neogastropoda: The Backbone Hidden in the Bush. Syst Biol. 73:521–531.

Field D.J., Berv J.S., Hsiang A.Y., Lanfear R., Landis M.J., Dornburg A. Timing the extant avian radiation: The rise of modern birds, and the importance of modeling molecular rate variation.

Fleming J.F., Valero-Gracia A., Struck T.H. 2023. Identifying and addressing methodological incongruence in phylogenomics: A review. Evol Appl. 16:1087–1104.

Foley N.M., Mason V.C., Harris A.J., Bredemeyer K.R., Damas J., Lewin H.A., Eizirik E., Gatesy J., Karlsson E.K., Lindblad-Toh K., Zoonomia Consortium‡, Springer M.S., Murphy W.J. 2023. A genomic timescale for placental mammal evolution. Science. 380:eabl8189.

Freitas F.V., Branstetter M.G., Griswold T., Almeida E.A.B. 2021. Partitioned Gene-Tree Analyses and Gene-Based Topology Testing Help Resolve Incongruence in a Phylogenomic Study of Host-Specialist Bees (Apidae: Eucerinae). Mol Biol Evol. 38:1090–1100.

Frumkin I., Lajoie M.J., Gregg C.J., Hornung G., Church G.M., Pilpel Y. 2018. Codon usage of highly expressed genes affects proteome-wide translation efficiency. Proc Natl Acad Sci U S A. 115:E4940–E4949.

Ghezelayagh A., Harrington R.C., Burress E.D., Campbell M.A., Buckner J.C., Chakrabarty P., Glass J.R., McCraney W.T., Unmack P.J., Thacker C.E., Alfaro M.E., Friedman S.T., Ludt W.B., Cowman P.F., Friedman M., Price S.A., Dornburg A., Faircloth B.C., Wainwright P.C., Near T.J. 2022. Prolonged morphological expansion of spiny-rayed fishes following the end-Cretaceous. Nat Ecol Evol. 6:1211–1220.

Gosselin S., Fullmer M.S., Feng Y., Gogarten J.P. 2022. Improving Phylogenies Based on Average Nucleotide Identity, Incorporating Saturation Correction and Nonparametric Bootstrap Support. Syst Biol. 71:396–409.

Grigoriadis K., Huebner A., Bunkum A., Colliver E., Frankell A.M., Hill M.S., Thol K., Birkbak N.J., Swanton C., Zaccaria S., McGranahan N. 2024. CONIPHER: a computational framework for scalable phylogenetic reconstruction with error correction. Nat Protoc. 19:159–183.

Haber M.H. 2025. Positively misleading errors. Synthese. 206.

Hanson G., Coller J. 2018. Codon optimality, bias and usage in translation and mRNA decay. Nat Rev Mol Cell Biol. 19:20–30.

Hendy M.D., Penny D. 1989. A Framework for the Quantitative Study of Evolutionary Trees. Systematic Zoology. 38:297.

Höhna S., Lower S.E., Duchen P., Catalán A. 2025. Robustness of divergence time estimation despite gene tree estimation error: a case study of fireflies (Coleoptera: Lampyridae). Syst Biol. 74:335–348.

Ho S.Y., Jermiin L. 2004. Tracing the decay of the historical signal in biological sequence data. Syst Biol. 53:623–637.

Huelsenbeck J.P. 1997. Is the Felsenstein Zone a Fly Trap? Systematic Biology. 46:69–74.

Jarvis E.D., Mirarab S., Aberer A.J., Li B., Houde P., Li C., Ho S.Y.W., Faircloth B.C., Nabholz B., Howard J.T., Suh A., Weber C.C., da Fonseca R.R., Li J., Zhang F., Li H., Zhou L., Narula N., Liu L., Ganapathy G., Boussau B., Bayzid M.S., Zavidovych V., Subramanian S., Gabaldón T., Capella-Gutiérrez S., Huerta-Cepas J., Rekepalli B., Munch K., Schierup M., Lindow B., Warren W.C., Ray D., Green R.E., Bruford M.W., Zhan X., Dixon A., Li S., Li N., Huang Y., Derryberry E.P., Bertelsen M.F., Sheldon F.H., Brumfield R.T., Mello C.V., Lovell P.V., Wirthlin M., Schneider M.P.C., Prosdocimi F., Samaniego J.A., Vargas Velazquez A.M., Alfaro-Núñez A., Campos P.F., Petersen B., Sicheritz-Ponten T., Pas A., Bailey T., Scofield P., Bunce M., Lambert D.M., Zhou Q., Perelman P., Driskell A.C., Shapiro B., Xiong Z., Zeng Y., Liu S., Li Z., Liu B., Wu K., Xiao J., Yinqi X., Zheng Q., Zhang Y., Yang H., Wang J., Smeds L., Rheindt F.E., Braun M., Fjeldsa J., Orlando L., Barker F.K., Jønsson K.A., Johnson W., Koepfli K.-P., O’Brien S., Haussler D., Ryder O.A., Rahbek C., Willerslev E., Graves G.R., Glenn T.C., McCormack J., Burt D., Ellegren H., Alström P., Edwards S.V., Stamatakis A., Mindell D.P., Cracraft J., Braun E.L., Warnow T., Jun W., Gilbert M.T.P., Zhang G. 2014. Whole-genome analyses resolve early branches in the tree of life of modern birds. Science. 346:1320–1331.

Jermiin L.S., Catullo R.A., Holland B.R. 2020. A new phylogenetic protocol: dealing with model misspecification and confirmation bias in molecular phylogenetics. NAR Genom Bioinform. 2:lqaa041.

Kapli P., Yang Z., Telford M.J. 2020. Phylogenetic tree building in the genomic age. Nat Rev Genet. 21:428–444.

Klopfstein S., Massingham T., Goldman N. 2017. More on the Best Evolutionary Rate for Phylogenetic Analysis. Syst Biol. 66:769–785.

Kuang T., Tornabene L., Li J., Jiang J., Chakrabarty P., Sparks J.S., Naylor G.J.P., Li C. 2018. Phylogenomic analysis on the exceptionally diverse fish clade Gobioidei (Actinopterygii: Gobiiformes) and data-filtering based on molecular clocklikeness. Mol Phylogenet Evol. 128:192–202.

Lanfear R., Hahn M.W. 2024. The Meaning and Measure of Concordance Factors in Phylogenomics. Mol Biol Evol. 41.

Lartillot N. 2013. Phylogenetic patterns of GC-biased gene conversion in placental mammals and the evolutionary dynamics of recombination landscapes. Mol. Biol. Evol. 30:489–502.

Lewis P.O., Chen M.-H., Kuo L., Lewis L.A., Fučíková K., Neupane S., Wang Y.-B., Shi D. 2016. Estimating Bayesian Phylogenetic Information Content. Syst. Biol.

Manuel C., Sakalli E., Schmidt H.A., Viñas C., von Haeseler A., Elgert C. 2025. When the Past Fades: Detecting Phylogenetic Signal with SatuTe. Mol Biol Evol. 42.

Martinez-Gutierrez C.A., Aylward F.O. 2021. Phylogenetic Signal, Congruence, and Uncertainty across Bacteria and Archaea. Mol Biol Evol. 38:5514–5527.

McCarthy C.G.P., Mulhair P.O., Siu-Ting K., Creevey C.J., O’Connell M.J. 2023. Improving Orthologous Signal and Model Fit in Datasets Addressing the Root of the Animal Phylogeny. Mol Biol Evol. 40.

McCraney W.T., Thacker C.E., Faircloth B.C., Harrington R.C., Near T.J., Alfaro M.E. 2025. Explosion of goby fish diversity at the Eocene-Oligocene transition. Mol Phylogenet Evol. 207:108342.

McTavish E.J., Steel M., Holder M.T. 2015. Twisted trees and inconsistency of tree estimation when gaps are treated as missing data - The impact of model mis-specification in distance corrections. Mol Phylogenet Evol. 93:289–295.

Minh B.Q., Hahn M.W., Lanfear R. 2020. New Methods to Calculate Concordance Factors for Phylogenomic Datasets. Mol Biol Evol. 37:2727–2733.

de Moya R.S., Yoshizawa K., Walden K.K.O., Sweet A.D., Dietrich C.H., Kevin P J. 2021. Phylogenomics of Parasitic and Nonparasitic Lice (Insecta: Psocodea): Combining Sequence Data and Exploring Compositional Bias Solutions in Next Generation Data Sets. Syst Biol. 70:719–738.

Mo Y.K., Lanfear R., Hahn M.W., Minh B.Q. 2023. Updated site concordance factors minimize effects of homoplasy and taxon sampling. Bioinformatics. 39.

Nagy L.G., Szöllősi G. 2017. Fungal Phylogeny in the Age of Genomics: Insights Into Phylogenetic Inference From Genome-Scale Datasets. Adv Genet. 100:49–72.

Naser-Khdour S., Minh B.Q., Lanfear R. 2026. Phylogenetic Accuracy Under Non-Stationary and Non-Homogeneous Conditions: A Simulation Study. Syst Biol.

Near T.J., Thacker C.E. 2024. Phylogenetic classification of living and fossil ray-finned fishes (Actinopterygii). Bull. Peabody Mus. Nat. Hist. 65.

Ontano A.Z., Gainett G., Aharon S., Ballesteros J.A., Benavides L.R., Corbett K.F., Gavish-Regev E., Harvey M.S., Monsma S., Santibáñez-López C.E., Setton E.V.W., Zehms J.T., Zeh J.A., Zeh D.W., Sharma P.P. 2021. Taxonomic Sampling and Rare Genomic Changes Overcome Long-Branch Attraction in the Phylogenetic Placement of Pseudoscorpions. Mol Biol Evol. 38:2446–2467.

Parker E., Dornburg A., Domínguez-Domínguez O., Piller K.R. 2019. Assessing phylogenetic information to reveal uncertainty in historical data: An example using Goodeinae (Teleostei: Cyprinodontiformes: Goodeidae). Mol. Phylogenet. Evol. 134:282–290.

Philippe H., Brinkmann H., Lavrov D.V., Littlewood D.T.J., Manuel M., Wörheide G., Baurain D. 2011. Resolving difficult phylogenetic questions: why more sequences are not enough. PLoS Biol. 9:e1000602.

Philippe H., Zhou Y., Brinkmann H., Rodrigue N., Delsuc F. 2005. Heterotachy and long-branch attraction in phylogenetics. BMC Evol. Biol. 5:50.

Prum R.O., Berv J.S., Dornburg A., Field D.J., Townsend J.P., Lemmon E.M., Lemmon A.R. 2015. A comprehensive phylogeny of birds (Aves) using targeted next-generation DNA sequencing. Nature. 526:569–573.

Quax T.E.F., Claassens N.J., Söll D., van der Oost J. 2015. Codon Bias as a Means to Fine-Tune Gene Expression. Mol Cell. 59:149–161.

Reddy S., Kimball R.T., Pandey A., Hosner P.A., Braun M.J., Hackett S.J., Han K.-L., Harshman J., Huddleston C.J., Kingston S., Marks B.D., Miglia K.J., Moore W.S., Sheldon F.H., Witt C.C., Yuri T., Braun E.L. 2017. Why Do Phylogenomic Data Sets Yield Conflicting Trees? Data Type Influences the Avian Tree of Life more than Taxon Sampling. Syst. Biol. 66:857–879.

Romiguier J., Cameron S.A., Woodard S.H., Fischman B.J., Keller L., Praz C.J. 2016. Phylogenomics Controlling for Base Compositional Bias Reveals a Single Origin of Eusociality in Corbiculate Bees. Mol. Biol. Evol. 33:670–678.

Romiguier J., Ranwez V., Delsuc F., Galtier N., Douzery E.J.P. 2013. Less is more in mammalian phylogenomics: AT-rich genes minimize tree conflicts and unravel the root of placental mammals. Mol. Biol. Evol. 30:2134–2144.

Romiguier J., Roux C. 2017. Analytical Biases Associated with GC-Content in Molecular Evolution. Front. Genet. 8:246001.

Rota J., Twort V., Chiocchio A., Peña C., Wheat C.W., Kaila L., Wahlberg N. 2022. The unresolved phylogenomic tree of butterflies and moths (Lepidoptera): Assessing the potential causes and consequences. Syst. Entomol. 47:531–550.

Rousselle M., Laverré A., Figuet E., Nabholz B., Galtier N. 2019. Influence of Recombination and GC-biased Gene Conversion on the Adaptive and Nonadaptive Substitution Rate in Mammals versus Birds. Mol. Biol. Evol. 36:458–471.

Salichos L., Leonidas S., Antonis R. 2013. Inferring ancient divergences requires genes with strong phylogenetic signals. Nature. 497:327–331.

Shen X.-X., Hittinger C.T., Rokas A. 2017. Contentious relationships in phylogenomic studies can be driven by a handful of genes. Nat Ecol Evol. 1:126.

Singhal S., Colston T.J., Grundler M.R., Smith S.A., Costa G.C., Colli G.R., Moritz C., Pyron R.A., Rabosky D.L. 2021. Congruence and Conflict in the Higher-Level Phylogenetics of Squamate Reptiles: An Expanded Phylogenomic Perspective. Syst Biol. 70:542–557.

Soltis P.S., Soltis D.E., Savolainen V., Crane P.R., Barraclough T.G. 2002. Rate heterogeneity among lineages of tracheophytes: integration of molecular and fossil data and evidence for molecular living fossils. Proc. Natl. Acad. Sci. U. S. A. 99:4430–4435.

Spinks P.Q., Thomson R.C., Lovely G.A., Shaffer H.B. 2009. Assessing what is needed to resolve a molecular phylogeny: simulations and empirical data from emydid turtles. BMC Evol Biol. 9:56.

Steenwyk J.L., Li Y., Zhou X., Shen X.-X., Rokas A. 2023. Incongruence in the phylogenomics era. Nat Rev Genet. 24:834–850.

Stewart A.A., Wiens J.J. 2025. A time-calibrated salamander phylogeny including 765 species and 503 genes. Mol Phylogenet Evol. 204:108272.

Stiller J., Feng S., Chowdhury A.-A., Rivas-González I., Duchêne D.A., Fang Q., Deng Y., Kozlov A., Stamatakis A., Claramunt S., Nguyen J.M.T., Ho S.Y.W., Faircloth B.C., Haag J., Houde P., Cracraft J., Balaban M., Mai U., Chen G., Gao R., Zhou C., Xie Y., Huang Z., Cao Z., Yan Z., Ogilvie H.A., Nakhleh L., Lindow B., Morel B., Fjeldså J., Hosner P.A., da Fonseca R.R., Petersen B., Tobias J.A., Székely T., Kennedy J.D., Reeve A.H., Liker A., Stervander M., Antunes A., Tietze D.T., Bertelsen M.F., Lei F., Rahbek C., Graves G.R., Schierup M.H., Warnow T., Braun E.L., Gilbert M.T.P., Jarvis E.D., Mirarab S., Zhang G. 2024. Complexity of avian evolution revealed by family-level genomes. Nature. 629:851–860.

Susko E. 2015. Bayesian long branch attraction bias and corrections. Syst Biol. 64:243–255.

Susko E., Roger A.J. 2021. Long Branch Attraction Biases in Phylogenetics. Syst Biol. 70:838–843.

Su Z., Townsend J.P. 2015. Utility of characters evolving at diverse rates of evolution to resolve quartet trees with unequal branch lengths: analytical predictions of long-branch effects. BMC Evol. Biol. 15:86.

Su Z., Zhuo S., Zheng W., Francesc L.-G., Townsend J.P. 2014. The impact of incorporating molecular evolutionary model into predictions of phylogenetic signal and noise. Frontiers in Ecology and Evolution. 2.

Szánthó L.L., Lartillot N., Szöllősi G.J., Schrempf D. 2023. Compositionally Constrained Sites Drive Long-Branch Attraction. Syst Biol. 72:767–780.

Torruella G., Galindo L.J., Moreira D., López-García P. 2025. Phylogenomics of neglected flagellated protists supports a revised eukaryotic tree of life. Curr Biol. 35:198–207.e4.

Townsend J.P. 2007. Profiling phylogenetic informativeness. Syst. Biol. 56:222–231.

Townsend J.P., Lopez-Giraldez F. 2010. Optimal selection of gene and ingroup taxon sampling for resolving phylogenetic relationships. Syst Biol. 59:446–457.

Townsend J.P., Su Z., Tekle Y.I. 2012. Phylogenetic signal and noise: predicting the power of a data set to resolve phylogeny. Syst. Biol. 61:835–849.

Vankan M., Ho S.Y.W., Duchêne D.A. 2022. Evolutionary Rate Variation among Lineages in Gene Trees has a Negative Impact on Species-Tree Inference. Syst Biol. 71:490–500.

Wang Y., Li Y.-D., Wang S., Tihelka E., Engel M.S., Cai C. 2025. Modeling compositional heterogeneity resolves deep phylogeny of flowering plants. Plant Divers. 47:13–20.

White N.D., Braun M.J. 2019. Extracting phylogenetic signal from phylogenomic data: Higher-level relationships of the nightbirds (Strisores). Mol. Phylogenet. Evol. 141:106611.

Williams T.A., Cox C.J., Foster P.G., Szöllősi G.J., Embley T.M. 2020. Phylogenomics provides robust support for a two-domains tree of life. Nat Ecol Evol. 4:138–147.

Wortley A.H., Rudall P.J., Harris D.J., Scotland R.W. 2005. How much data are needed to resolve a difficult phylogeny?: case study in Lamiales. Syst Biol. 54:697–709.

Wu X., Xu M., Yang J.-R., Lu J. 2024. Genome-wide impact of codon usage bias on translation optimization in Drosophila melanogaster. Nat Commun. 15:8329.

Xia X., Xie Z., Salemi M., Chen L., Wang Y. 2003. An index of substitution saturation and its application. Mol. Phylogenet. Evol. 26:1–7.

Yang Z. 1998. On the best evolutionary rate for phylogenetic analysis. Syst Biol. 47:125–133.

Young A.D., Gillung J.P. 2020. Phylogenomics — principles, opportunities and pitfalls of big-data phylogenetics. Syst. Entomol. 45:225–247.

Zhang Y.M., Williams J.L., Lucky A. 2019. Understanding UCEs: A comprehensive primer on using ultraconserved elements for arthropod phylogenomics. Insect Syst. Divers. 3.

Zhao M., Kurtis S.M., White N.D., Moncrieff A.E., Leite R.N., Brumfield R.T., Braun E.L., Kimball R.T. 2023. Exploring Conflicts in Whole Genome Phylogenetics: A Case Study Within Manakins (Aves: Pipridae). Syst Biol. 72:161–178.

Zhou Z., Dang Y., Zhou M., Li L., Yu C.-H., Fu J., Chen S., Liu Y. 2016. Codon usage is an important determinant of gene expression levels largely through its effects on transcription. Proc Natl Acad Sci U S A. 113:E6117–E6125.

Zuntini A.R., Carruthers T., Maurin O., Bailey P.C., Leempoel K., Brewer G.E., Epitawalage N., Françoso E., Gallego-Paramo B., McGinnie C., Negrão R., Roy S.R., Simpson L., Toledo Romero E., Barber V.M.A., Botigué L., Clarkson J.J., Cowan R.S., Dodsworth S., Johnson M.G., Kim J.T., Pokorny L., Wickett N.J., Antar G.M., DeBolt L., Gutierrez K., Hendriks K.P., Hoewener A., Hu A.-Q., Joyce E.M., Kikuchi I.A.B.S., Larridon I., Larson D.A., de Lírio E.J., Liu J.-X., Malakasi P., Przelomska N.A.S., Shah T., Viruel J., Allnutt T.R., Ameka G.K., Andrew R.L., Appelhans M.S., Arista M., Ariza M.J., Arroyo J., Arthan W., Bachelier J.B., Bailey C.D., Barnes H.F., Barrett M.D., Barrett R.L., Bayer R.J., Bayly M.J., Biffin E., Biggs N., Birch J.L., Bogarín D., Borosova R., Bowles A.M.C., Boyce P.C., Bramley G.L.C., Briggs M., Broadhurst L., Brown G.K., Bruhl J.J., Bruneau A., Buerki S., Burns E., Byrne M., Cable S., Calladine A., Callmander M.W., Cano Á., Cantrill D.J., Cardinal-McTeague W.M., Carlsen M.M., Carruthers A.J.A., de Castro Mateo A., Chase M.W., Chatrou L.W., Cheek M., Chen S., Christenhusz M.J.M., Christin P.-A., Clements M.A., Coffey S.C., Conran J.G., Cornejo X., Couvreur T.L.P., Cowie I.D., Csiba L., Darbyshire I., Davidse G., Davies N.M.J., Davis A.P., van Dijk K.-J., Downie S.R., Duretto M.F., Duvall M.R., Edwards S.L., Eggli U., Erkens R.H.J., Escudero M., de la Estrella M., Fabriani F., Fay M.F., Ferreira P. de L., Ficinski S.Z., Fowler R.M., Frisby S., Fu L., Fulcher T., Galbany-Casals M., Gardner E.M., German D.A., Giaretta A., Gibernau M., Gillespie L.J., González C.C., Goyder D.J., Graham S.W., Grall A., Green L., Gunn B.F., Gutiérrez D.G., Hackel J., Haevermans T., Haigh A., Hall J.C., Hall T., Harrison M.J., Hatt S.A., Hidalgo O., Hodkinson T.R., Holmes G.D., Hopkins H.C.F., Jackson C.J., James S.A., Jobson R.W., Kadereit G., Kahandawala I.M., Kainulainen K., Kato M., Kellogg E.A., King G.J., Klejevskaja B., Klitgaard B.B., Klopper R.R., Knapp S., Koch M.A., Leebens-Mack J.H., Lens F., Leon C.J., Léveillé-Bourret É., Lewis G.P., Li D.-Z., Li L., Liede-Schumann S., Livshultz T., Lorence D., Lu M., Lu-Irving P., Luber J., Lucas E.J., Luján M., Lum M., Macfarlane T.D., Magdalena C., Mansano V.F., Masters L.E., Mayo S.J., McColl K., McDonnell A.J., McDougall A.E., McLay T.G.B., McPherson H., Meneses R.I., Merckx V.S.F.T., Michelangeli F.A., Mitchell J.D., Monro A.K., Moore M.J., Mueller T.L., Mummenhoff K., Munzinger J., Muriel P., Murphy D.J., Nargar K., Nauheimer L., Nge F.J., Nyffeler R., Orejuela A., Ortiz E.M., Palazzesi L., Peixoto A.L., Pell S.K., Pellicer J., Penneys D.S., Perez-Escobar O.A., Persson C., Pignal M., Pillon Y., Pirani J.R., Plunkett G.M., Powell R.F., Prance G.T., Puglisi C., Qin M., Rabeler R.K., Rees P.E.J., Renner M., Roalson E.H., Rodda M., Rogers Z.S., Rokni S., Rutishauser R., de Salas M.F., Schaefer H., Schley R.J., Schmidt-Lebuhn A., Shapcott A., Al-Shehbaz I., Shepherd K.A., Simmons M.P., Simões A.O., Simões A.R.G., Siros M., Smidt E.C., Smith J.F., Snow N., Soltis D.E., Soltis P.S., Soreng R.J., Sothers C.A., Starr J.R., Stevens P.F., Straub S.C.K., Struwe L., Taylor J.M., Telford I.R.H., Thornhill A.H., Tooth I., Trias-Blasi A., Udovicic F., Utteridge T.M.A., Del Valle J.C., Verboom G.A., Vonow H.P., Vorontsova M.S., de Vos J.M., Al-Wattar N., Waycott M., Welker C.A.D., White A.J., Wieringa J.J., Williamson L.T., Wilson T.C., Wong S.Y., Woods L.A., Woods R., Worboys S., Xanthos M., Yang Y., Zhang Y.-X., Zhou M.-Y., Zmarzty S., Zuloaga F.O., Antonelli A., Bellot S., Crayn D.M., Grace O.M., Kersey P.J., Leitch I.J., Sauquet H., Smith S.A., Eiserhardt W.L., Forest F., Baker W.J. 2024. Phylogenomics and the rise of the angiosperms. Nature. 629:843–850.

