## Appendix for "Signal, noise, and bias in phylogenetic inference: potential and limits to the resolution of phylogenetic trees in the phylogenomic era"

### Calculation of Signal, Noise, and Bias for a Quartet

*Signal*

For a large number of characters $n$—typical for gene sequences—the unidirectional random walk distribution of $Y_3^{SUM}$, the cumulative number of characters that exhibit true synapomorphy asymptotically converges onto a continuous Gaussian distribution (Townsend et al. 2012), whose expected value and variance are, respectively,

$$\mathrm{E}[Y_3^{SUM}] = \mathrm{E}[\sum_{i=1}^{n} \Pi_i] = \sum_{i=1}^{n} \mathrm{E}[\Pi_i] = \sum_{i=1}^{n} \Pi(i) \qquad \textbf{(S1)}$$

and

$$\mathrm{VAR}[Y_3^{SUM}] = \mathrm{VAR}[\sum_{i=1}^{n} \Pi_i] = \sum_{i=1}^{n} \mathrm{VAR}[\Pi_i]$$

$$= \sum_{i=1}^{n} \mathrm{E}[\Pi_i^{\,2}] - \mathrm{E}[\Pi_i]^2$$

$$= \sum_{i=1}^{n} \Pi(i) \times 1^2 - \Pi(i)^2$$

$$= \sum_{i=1}^{n} \Pi(i)(1 - \Pi(i)) \,. \qquad \textbf{(S2)}$$

Evaluating **Equation S1** and **S2** thus yields the expected value and variance of signal, respectively.

*Noise*

Similarly, $X_1^{SUM}$, $X_2^{SUM}$, and $X_3^{SUM}$ all asymptotically converge onto a continuous Gaussian distribution assuming a large number of characters (Townsend et al. 2012). Their expected values and variances can be similarly evaluated and expressed as:

$$\mathrm{E}[X_1^{SUM}] = \sum_{i=1}^{n} x_1(i) \text{, and} \tag{S3}$$

$$\mathrm{VAR}[X_1^{SUM}] = \sum_{i=1}^{n} x_1(i)(1 - x_1(i)) \text{;} \tag{S4}$$

$$\mathrm{E}[X_2^{SUM}] = \sum_{i=1}^{n} x_2(i) \text{, and} \tag{S5}$$

$$\mathrm{VAR}[X_2^{SUM}] = \sum_{i=1}^{n} x_2(i)(1 - x_2(i)) \text{;} \tag{S6}$$

$$\mathrm{E}[X_3^{SUM}] = \sum_{i=1}^{n} x_3(i) \text{, and} \tag{S7}$$

$$\mathrm{VAR}[X_3^{SUM}] = \sum_{i=1}^{n} x_3(i)(1 - x_3(i)) \text{;} \tag{S8}$$

To evaluate the expected value and variance of the predicted support for incorrect quartet resolution, *W*, we rearrange Equation 2:

$$\begin{aligned} \mathrm{W} &= \mathrm{MAX}[X_1^{SUM}, X_2^{SUM}] - X_3^{SUM} \\ &= \mathrm{MAX}[X_1^{SUM} - X_3^{SUM}, X_2^{SUM} - X_3^{SUM}] \text{.} \end{aligned} \tag{S9}$$

Since $X_1^{SUM}$, $X_2^{SUM}$, and $X_3^{SUM}$ can all be approximated as Gaussian distributions, their differences, $X_1^{SUM} - X_3^{SUM}$ and $X_2^{SUM} - X_3^{SUM}$, are also approximately Gaussian. Denoting the expected values of $X_1^{SUM} - X_3^{SUM}$ and $X_2^{SUM} - X_3^{SUM}$ as $\mu_1$ and $\mu_2$, respectively, their variances as $\sigma_1^2$ and $\sigma_2^2$, and their correlation coefficient as $\rho$, from Nadarajah and Kotz (2008), the first two moments of the probability distribution of *W* are:

$$\mathrm{E}[W] = \mu_1 \Phi\left(\frac{\mu_1 - \mu_2}{\theta}\right) + \mu_2 \Phi\left(\frac{\mu_2 - \mu_1}{\theta}\right) + \theta\phi\left(\frac{\mu_1 - \mu_2}{\theta}\right), \text{ and} \quad \text{(S10)}$$

$$\mathrm{E}[W^2] = (\sigma_1^2 + \mu_1^2)\Phi\left(\frac{\mu_1 - \mu_2}{\theta}\right) + (\sigma_2^2 + \mu_2^2)\Phi\left(\frac{\mu_2 - \mu_1}{\theta}\right) + (\mu_1 + \mu_2)\theta\phi\left(\frac{\mu_1 - \mu_2}{\theta}\right). \quad \text{(S11)}$$

where $\phi(\cdot)$ and $\Phi(\cdot)$ are, respectively, the probability distribution function and the cumulative distribution function of the standard normal distribution, and additionally

$$\theta = \sqrt{\sigma_1^2 + \sigma_2^2 - 2\rho\sigma_1\sigma_2}. \quad \text{(S12)}$$

The variance of *W* is thus given by:

$$\mathrm{VAR}[W] = \mathrm{E}[W^2] - \left(\mathrm{E}[W]\right)^2. \quad \text{(S13)}$$

From Equations S3, S5, and S7,

$$\mu_1 = \mathrm{E}[X_1^{SUM} - X_3^{SUM}] = \sum_{i=1}^{n} x_1(i) - x_3(i), \text{ and} \quad \text{(S14)}$$

$$\mu_2 = \mathrm{E}[X_2^{SUM} - X_3^{SUM}] = \sum_{i=1}^{n} x_2(i) - x_3(i) \quad . \tag{S15}$$

We expand the expression for the variance $\sigma_1^2$ :

$$\sigma_1^2 = \mathrm{VAR}[X_1^{SUM} - X_3^{SUM}] = \mathrm{VAR}[X_1^{SUM}] + \mathrm{VAR}[X_3^{SUM}] - 2\mathrm{Cov}[X_1^{SUM}, X_3^{SUM}] \, . \tag{S16}$$

For $\mathrm{Cov}(X_1^{SUM}, X_3^{SUM})$ , because $X_{1j}$ and $X_{3i}$ are independent for different characters (i.e. when $i \neq j$ ), the covariance terms are zero when $i \neq j$ . Then, by the bi-linearity property of covariance,

$$\mathrm{Cov}(X_1^{SUM}, X_3^{SUM}) = \mathrm{Cov}(\sum_{j=1}^{n} X_{1j}, \sum_{i=1}^{n} X_{3i}) = \sum_{i=1}^{n}\sum_{j=1}^{n} \mathrm{Cov}(X_{1j}, X_{3i})$$

$$= \sum_{i=1}^{n} \mathrm{Cov}(X_{1i}, X_{3i}) = \sum_{i=1}^{n} \mathrm{E}[X_{1i}X_{3i}] - \mathrm{E}[X_{1i}]\mathrm{E}[X_{3i}]$$

$$= \sum_{i=1}^{n} 0 - x_1(i)x_3(i) = \sum_{i=1}^{n} -x_1(i)x_3(i) \quad . \tag{S17}$$

Substituting Equations S4, S8, and S17 into Equation S16,

$$\sigma_1^2 = \sum_{i=1}^{n} x_1(i) + x_3(i) - (x_1(i) - x_3(i))^2 \quad . \tag{S18}$$

Therefore,

$$\sigma_1 = \sqrt{\sum_{i=1}^{n} x_1(i) + x_3(i) - (x_1(i) - x_3(i))^2} \quad . \tag{S19}$$

Following the same calculation as in Equations S16–19 for $\sigma_1$ ,

$$\sigma_2 = \sqrt{\sum_{i=1}^{n} x_2(i) + x_3(i) - (x_2(i) - x_3(i))^2} \quad \text{(S20)}$$

By definition, $\rho = \dfrac{\mathrm{Cov}(X_1^{SUM} - X_3^{SUM}, X_2^{SUM} - X_3^{SUM})}{\sigma_1 \sigma_2}$ . The numerator can be evaluated as

$$\begin{aligned}
&\mathrm{Cov}(X_1^{SUM} - X_3^{SUM}, X_2^{SUM} - X_3^{SUM}) \\
&= \mathrm{Cov}(X_1^{SUM}, X_2^{SUM}) - \mathrm{Cov}(X_1^{SUM}, X_3^{SUM}) - \mathrm{Cov}(X_3^{SUM}, X_2^{SUM}) + \mathrm{Cov}(X_3^{SUM}, X_3^{SUM}) \\
&= \mathrm{Cov}(X_1^{SUM}, X_2^{SUM}) - \mathrm{Cov}(X_1^{SUM}, X_3^{SUM}) - \mathrm{Cov}(X_2^{SUM}, X_3^{SUM}) + \mathrm{VAR}(X_3^{SUM}) .
\end{aligned}$$

(S21)

Using a similar derivation to Equation S17 for $\mathrm{Cov}(X_1^{SUM}, X_3^{SUM})$ , we obtain:

$$\mathrm{Cov}(X_1^{SUM}, X_2^{SUM}) = \sum_{i=1}^{n} \mathrm{Cov}(X_{1i}, X_{2i}) = \sum_{i=1}^{n} -x_1(i)x_2(i) \text{, and} \quad \text{(S22)}$$

$$\mathrm{Cov}(X_2^{SUM}, X_3^{SUM}) = \sum_{i=1}^{n} \mathrm{Cov}(X_{2i}, X_{3i}) = \sum_{i=1}^{n} -x_2(i)x_3(i) . \quad \text{(S23)}$$

Substituting Equations S8, S17, S22, and S23 into Equation S21 yields

$$\begin{aligned}
&\mathrm{Cov}(X_1^{SUM} - X_3^{SUM}, X_2^{SUM} - X_3^{SUM}) \\
&= \sum_{i=1}^{n} - x_1(i)x_2(i) + x_1(i)x_3(i) + x_2(i)x_3(i) + x_3(i)(1 - x_3(i)) .
\end{aligned} \quad \text{(S24)}$$

Substituting Equations S19, S20, and S24 into the definition for $\rho$ ,

$$\rho = \frac{\sum_{i=1}^{n} - x_1(i)x_2(i) + x_1(i)x_3(i) + x_2(i)x_3(i) + x_3(i)(1 - x_3(i))}{\sqrt{\sum_{i=1}^{n} x_1(i) + x_3(i) - (x_1(i) - x_3(i))^2} \sqrt{\sum_{i=1}^{n} x_2(i) + x_3(i) - (x_2(i) - x_3(i))^2}} \quad \text{(S25)}$$

Substituting Equations S19, S20, and S25 into Equation S12,

$$\theta = \sqrt{\sum_{i=1}^{n} x_1(i) + x_2(i) - (x_1(i) - x_2(i))^2} \quad \text{(S26)}$$

Then, the expected value and variance of $W$ can be obtained by evaluating Equations S10 and S13, respectively, using the numerically evaluated values of $\mu_1$ (Equations S14), $\mu_2$ (Equation S15), $\sigma_1^2$ (Equation S19), $\sigma_2^2$ (Equation S20), and $\rho$ (Equation S25). The expected value and variance of $W^*$ (*i.e.* noise) can be similarly evaluated under the alternative assumption of invariant nucleotide base frequencies over the quartet branches.

*Bias*

The expected value of bias, B, can be evaluated as the difference in the expected value of $W$ and the expected value of $W^*$ (*i.e.* noise):

$$\mathrm{E}[\mathrm{B}] = \mathrm{E}[W] - \mathrm{E}[W^*]. \quad \text{(S26)}$$

The variance of bias can be calculated as

$$VAR[B] = \int (t - E[B])^2 f(t)\, \mathrm{d}t \quad \text{(S27)}$$

where $f(t)$ is the probability distribution function of bias and is given by the probability distribution function of $W$ minus the probability distribution function of $W^*$.

From Nadarajah and Kotz (2008), the probability distribution function of $W$ is $g(t) = g_1(-t) + g_2(-t)$, where

$$g_1(t) = \frac{1}{\sigma_1}\phi\left(\frac{t+\mu_1}{\sigma_1}\right) \times \Phi\left(\frac{\rho(t+\mu_1)}{\sigma_1\sqrt{1-\rho^2}} - \frac{t+\mu_2}{\sigma_2\sqrt{1-\rho^2}}\right), \text{ and} \tag{S28}$$

$$g_2(t) = \frac{1}{\sigma_2}\phi\left(\frac{t+\mu_2}{\sigma_2}\right) \times \Phi\left(\frac{\rho(t+\mu_2)}{\sigma_2\sqrt{1-\rho^2}} - \frac{t+\mu_1}{\sigma_1\sqrt{1-\rho^2}}\right). \tag{S29}$$

Therefore, the probability distribution function of $W$ can be obtained by substituting the values of $\mu_1$, $\mu_2$, $\sigma_1^2$, $\sigma_2^2$, and $\rho$ into Equations S28 and S29. The probability distribution function of $W^*$ can be similarly evaluated under the alternative assumption of invariant nucleotide base frequencies over the quartet branches. Then, the variance of bias can be obtained by evaluating Equation S27.

*Signal, Noise, and Bias for a Hypothetical Quartet*

We now illustrate how signal, noise, and bias varies with respect to the data size (i.e. number of characters) by considering a hypothetical quartet tree with an internode of length $t_0 = 0.1$ (arbitrary time unit), and four even subtending branches $T_1 = T_2 = T_3 = T_4 = 1$. For simplicity, we assume that all characters in the data set evolve at the same rate $\lambda = 0.5$ (per time unit). From Equation S1, the expected value of signal increases linearly with the number of characters in the data set:

$$\mathrm{E}[Y_3^{SUM}] = \sum_{i=1}^{n} \Pi(i) \quad . \tag{S30}$$

The expected value of noise, $W^*$, can be evaluated via Equation S10 by assuming that nucleotide base frequencies are the same across the internode and subtending branches of the quartet tree. We use the superscript * to denote variables evaluated under this assumption of invariant nucleotide base across quartet branches. Now, since the four subtending branches of the quartet tree are even and symmetrical, support for either of the two incorrect quartet topologies is equal. In other words, $X_1^{SUM*}$ and $X_2^{SUM*}$ are equal, and so are $x_1^*(i)$ and $x_2^*(i)$. Equating $X_1^{SUM*}$ and $X_2^{SUM*}$ in Equations S4 and S5 leads to $\mu_1^* = \mu_2^*$. Equation S10 for $W^*$ can therefore be simplified:

$$\mathrm{E}[W^*] = \mu_1^* \Phi\left(\frac{\mu_1^* - \mu_2^*}{\theta^*}\right) + \mu_2^* \Phi\left(\frac{\mu_2^* - \mu_1^*}{\theta^*}\right) + \theta^* \phi\left(\frac{\mu_1^* - \mu_2^*}{\theta^*}\right)$$

$$= \mu_1^* \Phi(0) + \mu_2^* \Phi(0) + \theta^* \phi(0) = 2\mu_1^* \Phi(0) + \theta^* \phi(0)$$

$$= \sum_{i=1}^{n} x_1^*(i) - x_3^*(i) + \sqrt{\frac{\sum_{i=1}^{n} x_1^*(i) + x_2^*(i) - (x_1^*(i) - x_2^*(i))^2}{2\pi}}$$

$$= \sum_{i=1}^{n} x_1^*(i) - x_3^*(i) + \sqrt{\frac{\sum_{i=1}^{n} x_1^*(i)}{\pi}} \quad . \tag{S31}$$
